# Genetic diversity in global chicken breeds as a function of genetic distance to the wild populations

**DOI:** 10.1101/2020.01.29.924696

**Authors:** Dorcus Kholofelo Malomane, Steffen Weigend, Armin Otto Schmitt, Annett Weigend, Christian Reimer, Henner Simianer

## Abstract

Migration of populations from their founder population is expected to cause a reduction in genetic diversity and facilitates population differentiation between the populations and their founder population as predicted by the theory of genetic isolation by distance. Consistent with that, a model of expansion from a single founder predicts that patterns of genetic diversity in populations can be well explained by their geographic expansion from the founders, which is correlated to the genetic differentiation. To investigate this in the chicken, we have estimated the relationship between the genetic diversity in 172 domesticated chicken populations and their genetic distances to wild populations. We have found a strong inverse relationship whereby 87.5% of the variation in the overall genetic diversity of domesticated chicken can be explained by the genetic distance to the wild populations. We also investigated if different types of SNPs and genes present similar patterns of genetic diversity as the overall genome. Among different SNP classes, the non-synonymous ones were the most deviating from the overall genome. However, the genetic distances to wild populations still explained more variation in domesticated chicken diversity in all SNP classes ranging from 81.7 to 88.7%. The genetic diversity seemed to change at a faster rate within the chicken in genes that are associated with transmembrane transport, protein transport and protein metabolic processes, and lipid metabolic processes. In general, such genes are flexible to be manipulated according to the population needs. On the other hand, genes which the genetic diversity hardly changes despite the genetic distance to the wild populations are associated with major functions e.g. brain development. Therefore, changes in the genes may be detrimental to the chickens. These results contribute to the knowledge of different evolutionary patterns of different functional genomic regions in the chicken.

**Author summary:** The chicken was first domesticated about 6000 B.C. in Asia from the jungle fowl. Following domestication, chickens were taken to different parts of the world mainly by humans. Evolutionary forces such as selection and genetic drift have shaped diversification within the chicken species. In addition, new breeds or strains have been developed from crossbreeding programs facilitated by man. These events, together with other breeding practices, have led to genomic alterations causing genetic differentiation between the domesticated chickens and their ancestral/wild population as well as manipulation of the genetic diversity within the domesticated chickens. We investigated the relationship between 172 domesticated chicken populations from different selection, breeding and management backgrounds and their genetic distance to the wild type chickens. We found that the genetic diversity within the populations decreases with the increasing genetic distances to the wild types. Human manipulation of chicken genetic diversity has more effect on the genetic differentiation than simple geographic separations (through migrations) do. We further found that some genes associated with vital functions show evolutionary constraints or persistent selection across the populations and do not comply with this relationship i.e. the genetic diversity within the populations is constant despite the change in the genetic distance to the wild types.

## Introduction

Domesticated chickens (*Gallus gallus domesticus*) are one of the most widely distributed domestic animal species in the world. Some of the reasons are due to their portability and flexibility of transportation through human migration, stock trading, and expansion in the agricultural practices [1, 2], in addition their use for nutrition is not suffering from any religious or cultural reservations. It is commonly accepted that the world-spread chickens of today originate predominantly from domestication of the red jungle fowl (*Gallus gallus* species) in Asia (reviewed by Tixier-Boichard et al [3]). From the centers of domestication, chickens have dispersed into different parts of the world. There has been formation of new breeds or lines as populations moved outward from ancestral territories and settled in new colonies. One of the expectations from such expansion processes is the increase of genetic distances (increased differentiation) of the outward populations to the original ancestors, and the loss of genetic diversity within such populations due to genetic drift and subsequent serial founder effects [4–6]. In Malomane et al [7] we studied the overall genetic diversity between and within the chicken breeds. In the current study we aimed at investigating if the observed genetic diversity in the chicken breeds is a result of their genetic expansion from the chicken wild populations following the concepts behind the theory of genetic isolation by distance [8–10] and the model of expansion from a single location such as the ‘Out of Africa’ migration model [4]. The theory of genetic isolation by distance refers to the population genetic patterns whereby genetic differentiation increases with the increase in geographic distance between populations. This is because the exchange of genetic material between the populations (i.e. mating opportunities) is confined by the distance [8, 11]. Likewise, movements of individuals further apart from their founders would be expected to increase genetic differentiation. This has been established with the ‘Out of Africa’ theory which asserts that modern humans originate from Africa [13] and human populations worldwide resulted in a reduction in genetic diversity with the increasing geographic distance from east Africa (Ethiopia) [4, 5, 14, 15]. Similar studies in cattle also reported a decreasing genetic diversity with increasing geographic distance to the cattle domestication center in Southwest Asia [16, 17].

The loss of genetic diversity within the migrated populations, which can be explained by the geographic distance from their founders, is believed to be a good measure of neutral genetic diversity as a consequence of genetic drift. However, the overall genetic diversity is also a result of population specific events such as mutations, natural selection to favor adaptation in the current environments and/or artificial selection (e.g. in livestock production practices) as well as population specific drift [5]. Consequences of selection are often measured by non-neutral genetic variation as it is assumed that non-neutral regions with functional fitness effects in the genome evolve differently to the neutral genome. In this study we used the global collection of chicken breeds [7] to investigate the pattern of the overall genetic diversity moving outwards the centers of chicken domestication, given all events taking place in the genome. Furthermore, we investigate if different functional regions of the genome present similar patterns as the overall genome. We hypothesized that changes in genetic diversity may be faster in some genes or functional categories depending on their functions and changes may also be different in different breeds or breed groups due to different adaptive or artificial selection targets. Therefore, the pattern of relationship between genetic diversity and genetic distance may behave differently, less complying with the overall genome and more dynamic than the non-genic regions due to differences in selection patterns in addition to other population specific events.

Studying the theory of genetic isolation by distance and/or the concept of migration from a single location with chickens poses some challenges because the physical locations do not always represent their geographic origin (following migration from founders). For many chicken breeds the time point when they have migrated to their current locations is unknown. We also believe that geographic distances may not be the best predictor of the genetic diversity in the chicken. This is because unlike in humans where genetic evolution is mostly driven by natural circumstances, rapid migration, crossbreeding forced by man, refined breeding programs and artificial selection for desired traits have largely shaped the evolution of domesticated chickens. The changes in genetic diversity and evolutionary rates are often rapid in domesticated livestock and the genetic architecture of chickens around the same geographic location may also differ greatly depending on different breeding practices or selection targets. Therefore, in our study we used Reynolds’ genetic distances [19] instead of geographic distances but following similar concepts as the genetic isolation by distance and model of expansion from a single founder [5, 8, 9]. Reynolds’ distances estimate differences under the assumptions that genetic differentiation occurs by genetic drift.

## Results and discussion

### The relationship between the overall genetic diversity and the genetic distance to wild populations

The relationship between the observed heterozygosity within domestic chicken (*Gallus gallus domesticus*) populations and the genetic distance to the wild populations (*Gallus gallus*) is shown in Fig 1. The different breed categories as described in the Materials and Methods section and S1 Table are represented by symbols of different colours and shapes. There is a strong inverse relationship between the genetic diversity within populations and their genetic distances to the wild populations. This relationship is similar even when using just neutral markers (intergenic SNPs, Fig 2). Across these chicken populations, 87.5% (Table 1) of the total variation in the heterozygosity can be explained by the genetic distance to the wild populations. This figure is slightly higher than those obtained in several human studies when using geographic distances. Geographic distances of humans out of Africa explained 76.3% of microsatellite heterozygosity and 78.4% of fixation index *F*_*ST*_ variation in [5] and 85% of microsatellite heterozygosity in [15]. They had a correlation of −0.910 with SNP haplotype heterozygosity and −0.870 with microsatellite heterozygosity in the same study [20]. Furthermore, studies in humans have shown that there is a high correlation (e.g. 0.765 to 0.885 [5]) between the genetic distances (using different genetic distance measures) and geographic distance. However the correlations were not as high in domesticated cattle studies compared to humans. For example, a correlation of 0.624 was reported by [21] and while [16] reported a correlation of 0.750 for ancient cattle samples, the correlation was 0.540 in modern cattle samples. The weakening relationship between geographic and genetic distances in modern domesticated cattle was suggested to be due to the human manipulation of genetic diversity among other reasons, as it is with many domesticated livestock [16].

**Table 1.**
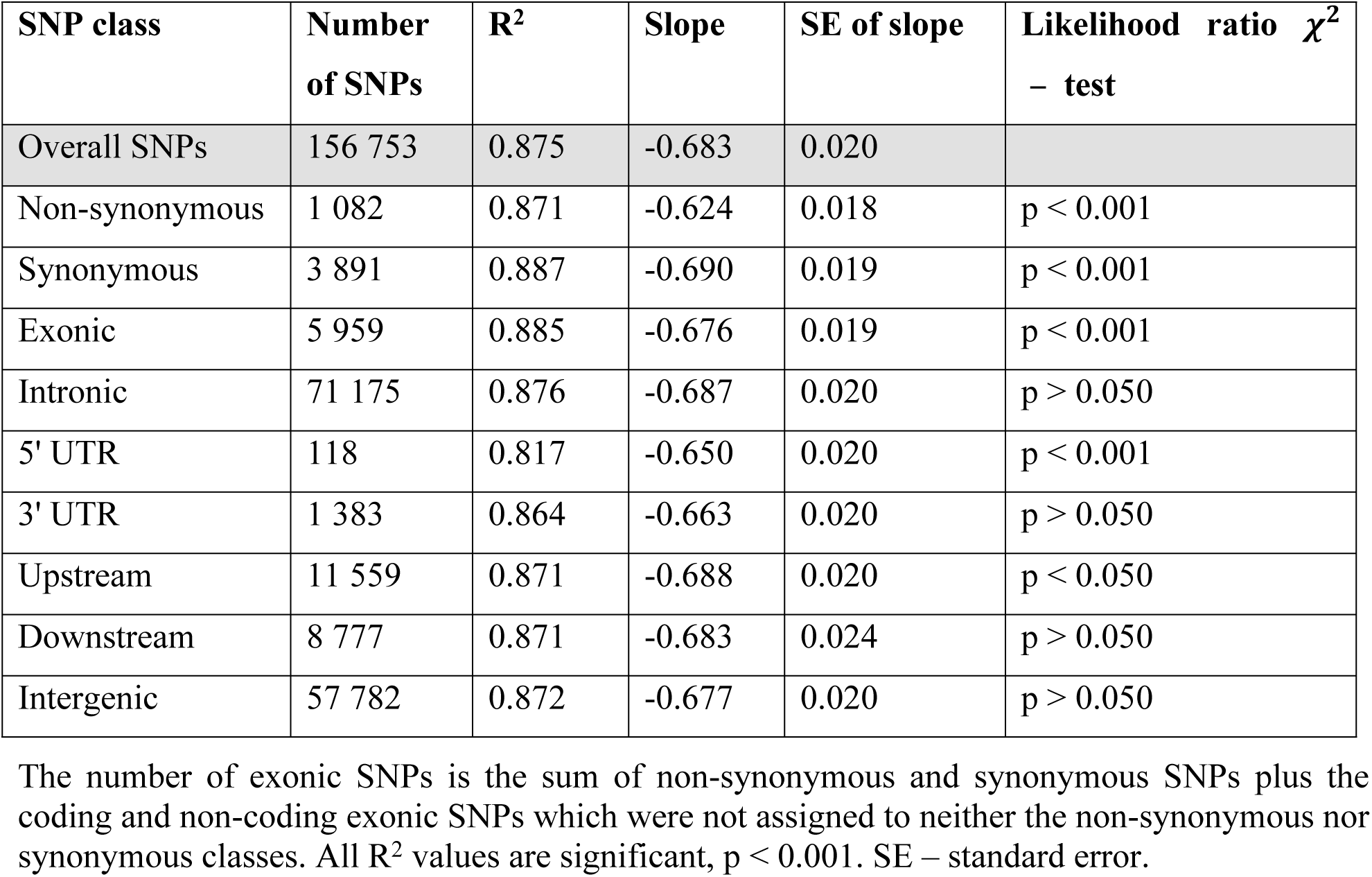
Comparisons of the linear relationship between genetic diversity and genetic distances of populations to *Gallus gallus* ssp. for different SNP classes.

| SNP class | Number of SNPs | R<sup>2</sup> | Slope | SE of slope | Likelihood ratio $\chi^2$ – test |
| --- | --- | --- | --- | --- | --- |
| Overall SNPs | 156 753 | 0.875 | -0.683 | 0.020 |  |
| Non-synonymous | 1 082 | 0.871 | -0.624 | 0.018 | p < 0.001 |
| Synonymous | 3 891 | 0.887 | -0.690 | 0.019 | p < 0.001 |
| Exonic | 5 959 | 0.885 | -0.676 | 0.019 | p < 0.001 |
| Intronic | 71 175 | 0.876 | -0.687 | 0.020 | p > 0.050 |
| 5' UTR | 118 | 0.817 | -0.650 | 0.020 | p < 0.001 |
| 3' UTR | 1 383 | 0.864 | -0.663 | 0.020 | p > 0.050 |
| Upstream | 11 559 | 0.871 | -0.688 | 0.020 | p < 0.050 |
| Downstream | 8 777 | 0.871 | -0.683 | 0.024 | p > 0.050 |
| Intergenic | 57 782 | 0.872 | -0.677 | 0.020 | p > 0.050 |

The number of exonic SNPs is the sum of non-synonymous and synonymous SNPs plus the coding and non-coding exonic SNPs which were not assigned to neither the non-synonymous nor synonymous classes. All R<sup>2</sup> values are significant, p < 0.001. SE – standard error.

**Fig 1.**
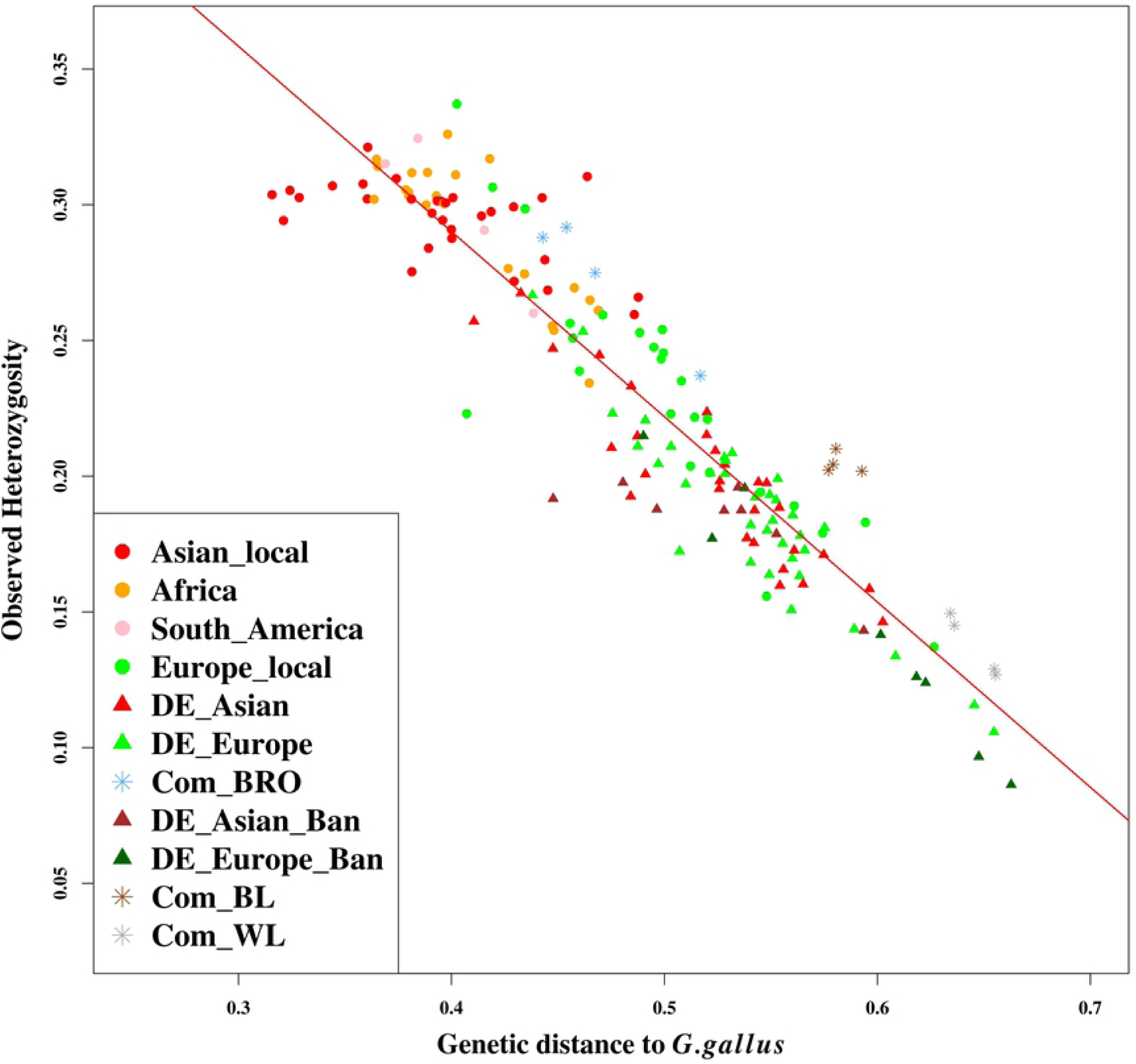
The relationship between the overall genetic diversity within populations and their genetic distance to *Gallus gallus*. The full names of the categories and description can be found in the S1 Table. The fitted regression line to the data with the equation heterozygosity = 0.563 – 0.683 x (genetic distance to *G. gallus*) is drawn in red. The R^2^ for the linear regression is 0.875 (p < 0.001).

**Fig 2.**
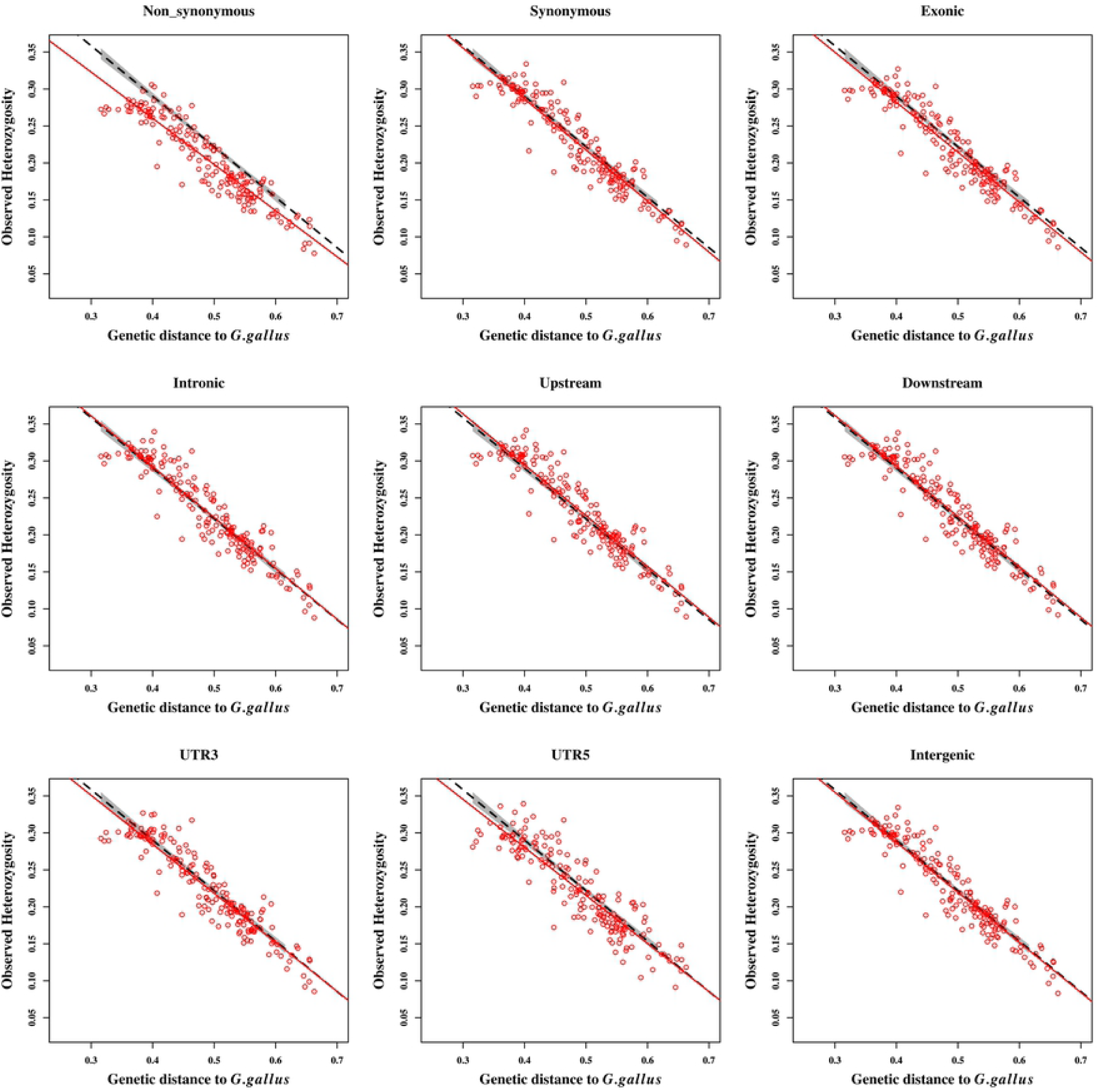
Genetic diversity within populations estimated from different SNP classes vs. their Reynolds’ genetic distance to *Gallus gallus* ssp. The red circles represent the 172 domesticated populations for the corresponding SNP class. Dashed black lines represent the regression lines for the relationship between observed heterozygosity and the genetic distance to *G. gallus* for the overall pattern and the red lines are for the SNP classes. The areas shaded in gray represent a 95% confidence interval. The R^2^ values and slopes of the linear relationships are shown in Table 1. UTR5 and UTR3 refer to the 5’ and 3’ UTR classes, respectively.

Since we had different population sizes whereby some population samples consisted of less than 15 individuals, we checked if this affects the estimates. We estimated the genetic diversity when only populations with 15 or more individuals were considered and found that the population sizes did not affect the estimates. We also sampled 1000 SNPs in 100 replicates to validate that the relationship between heterozygosity and genetic distance does not happen by chance. The percentages of variation explained in the 100 replicates ranged from 85.1 to 88.3% with a mean of 86.7%. S1 Fig shows the regression plots of the 100 replicates with their 95% confidence intervals. Furthermore, we permuted the SNPs to investigate whether the decreasing heterozygosity is not generally an artefact of the Reynolds’ distances. We found that the relationship between the observed heterozygosity and the genetic distance based on permutated SNPs was almost non-existing with an R2 value of 0.01. We also used the fixation index (*F*_*ST*_) as an alternative measure of differentiation, and found the Mantel correlation coefficient (r_m_) of pairwise *F*_*ST*_ values with the corresponding Reynolds’ distances to be 0.976. Reynolds’ genetic distances to the wild populations (*G .gallus*) and the *F*_*ST*_ values were highly correlated with a Pearson’s r = 0.990 and their relationship is shown in S2 Fig with an R^2^ value of 0.990. When using *F*_*ST*_, the genetic differentiation of the breeds from the wild populations (*G. gallus*) explained 86.2% of the variation in genetic diversity (S2 Fig).

Given our results we can conclude that the variance in genetic diversity within the domesticated chicken populations can be well explained by the genetic distance to the *Gallus gallus*. Although our current study may not directly prove this due to lack of geographic sampling coordinates, given the whole data set it is evident that the geographic distance alone may not well predict the observed genetic variations in the chickens because:

i. breeds of the same geographic origin are found scattered across the genetic diversity spectrum. This is the case for Asian (red symbols) and European (green symbols) type breeds. As it is shown in Fig 1 and as well highlighted in [7], the Asian and European chickens sampled from the German fancy breeders (denoted with prefix DE_) have highly reduced genetic diversity as well as higher genetic distance to the wild chickens (*G .gallus*) than their respective local breeds. However, when considering the sampling areas, the genetic diversity may correlates to the geographic distances to the *G. gallus* within the Asian breed categories but not in the European breeds. Many of the fancy breeds presumably originate from a small number of breeding birds imported from Asia to Europe. Following that, they have been subjected to strong phenotypic selection, with small effective population sizes, population bottlenecks, and intended inbreeding to keep the desired traits. Therefore, such practices are responsible for most of the variations in the genetic diversity of the fancy Asian and European type breeds vs. the respective local types.
ii. the concept of isolation by distance assumes that individuals from nearby locations are likely to be related due to mating possibilities. This is often the case in traditional breeding systems but it is not the case with the fancy and commercial breeding and management practices. Individuals within a commercial breeding herd are more related to each other than to other lines despite the geographic distances. In fancy breeds, there may be gene flow between small stocks based on personal contacts or personal relationships of breeders, but not related to geographic distance forming a substructure within the breed. Actually such gene flow between fancy breeds is also very limited. Furthermore, if geographic distance was a better predictor for the loss of genetic diversity and increased differentiation of breeds to the wild populations, then the African and South American breeds might be expected to have highly reduced genetic diversity due to geographic distances. They also would be expected to have high genetic distances to the wild populations as well as to the rest of the Asian populations; in fact, both expectations are not fulfilled, and some of the African populations were found to be clustered with the wild type breeds [7].

Therefore, the observed variations in genetic diversity may not well be predicted only by geographic expansion but rather by a combination with other aspects or subsequent events e.g. effective population sizes, types of breeding practices, and possibly subsequent series of founder events following the geographic expansion, as previously suggested [5, 6]. Such events which have taken place after geographic expansion have definitely contributed to the variations in allele frequencies and thus the genetic distances of domestic chickens to the wild populations. In addition, equilibrium between genetic drift, migration and mutation has probably not been reached in all studied populations, which would be compatible with the theory of genetic isolation by distance [5, 8, 9]. The theoretical expansion models are also based on ‘natural’ expansion through migration, while chickens and other livestock were actively transported by humans (e.g. with ships) to distant places.

### Comparisons of the patterns of genetic diversity between the overall genome (all SNPs) and different functional SNP classes

We compared the patterns of the relationship between the genetic diversity and genetic distances to the *Gallus gallus* species when using the overall SNPs to that from different SNP classes as shown in Fig 2 and Table 1. The rate of change in genetic diversity due to the genetic distance to the wild populations is represented by the slope in column 4. Compared to other SNP classes, the non-synonymous class showed a relevant deviation from the overall pattern whereby the observed heterozygosity across the breeds was lower than that of the overall genome. The non-synonymous class also had the most deviating slope among the classes (−0.624 compared to −0.683 for all SNPs). To investigate if the different pattern in the non-synonymous class is not due to the sample size, we resampled the same number of SNPs as in the non-synonymous class (1 082 SNPs) from the overall set (156K SNPs) 100 times. We estimated the heterozygosity and plotted the 100 samples to compare with the non-synonymous set. It is shown in S3 Fig that the difference in pattern of the non-synonymous class to the overall genome pattern is not due to the sample size.

Furthermore, the intergenic and intronic classes had the highest proportion of SNPs than the other SNP classes (Table 1). In order to validate that the similarity of these two classes to the overall is not an artefact of the sample sizes, we sampled 1000 SNPs a 100 times from the intergenic and intronic classes (separately). Then we estimated the heterozygosity and compared the results to the overall SNPs, showing that the similarities are not due to the larger sample sizes (S4 and S5 Figs). In comparing the regression models using the likelihood ratio test, the exonic (including both the synonymous and non-synonymous separately) and 5’ UTR SNP classes showed highly significant differences to the overall SNPs (p < 0.001, Table 1 last coloumn). Nonetheless, all SNP classes show a reduction in genetic diversity across populations with the increase in genetic distance to the wild types, with the R^2^ values ranging from 81.7% to 88.7%. The results show that for the synonymous SNPs, 88.7% of the variation in the heterozygosity across populations can be explained by their genetic distance to *G. gallus* while in the non-synonymous sites it explains 87.1%, and the lowest percentage was observed for 5’ UTR (81.7%). However, it is important to note that the 5’ UTR class had only 118 SNPs and hence the differences could be an effect of the sample size. To test this, we have randomly sampled 118 SNPs in 100 replicates from the overall set and estimated the relationship as we have done with the non-synonymous SNPs, and the R^2^ of the replicates ranged from 77.9% to 86.5% with a mean of 81.7%, suggesting that this result is most likely an artefact caused by small sample size.

Fig 3 shows the mean observed heterozygosity in the different SNP classes. Generally, the observed heterozygosity was lower in genic than in non-genic SNP classes. Within the genic class, lower heterozygosity was observed in exonic than in intronic SNPs. Consistent with Fig 2, the non-synonymous SNPs presented the lowest genetic diversity among all the SNP classes. This could be expected since non-synonymous changes can present favourable or disadvantagous consequences. The theoretical assumption is that selection acts rapidly towards fixation of the favourable alleles and purging of the non-favourable ones, thus leading to more homozygosity in these protein altering variants. The exonic and 5’ UTR classes followed the non-synonymous class with lowest mean heterozygosity. UTR variants can play a role in the regulation of gene expression and translation. For example, 3’ UTR could interfere with microRNA to facilitate the translation of critical disease genes (e.g. cancer genes in humans) [22, 23]. It is also claimed that positive selection for the adaptation of humans in different habitats has been achived with high differentiation in the 5’ UTR gene variants [24]. Such examples highlight the importance of UTR variants as possible targets for selection.

**Fig 3.**
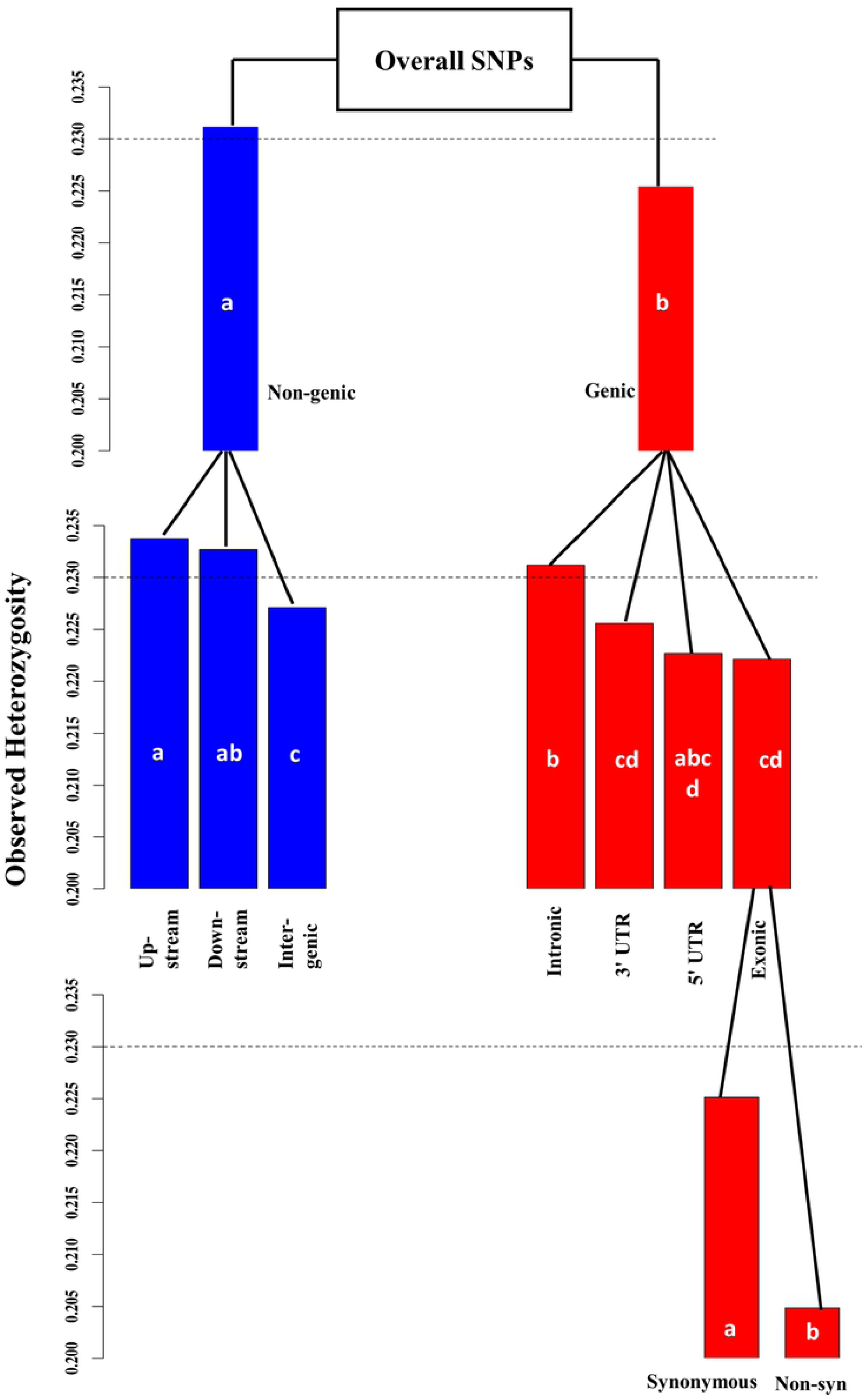
The mean observed heterozygosity in the different SNP classes. The gray dotted lines represent the overall mean observed heterozygosity when all SNPs are considered. Non-syn – Non-synonymous. The mean heterozygosities of the SNP classes were significantly different to the overall mean (Welch two sample t-test p < 0.05) except for the 3’ UTR and 5’ UTR classes. The standard errors (SEs) of the means were lower than 0.005 in all the SNP classes and the overall except for the 5’ UTR with SE = 0.009. Different letters in the bars means that there is significant difference in the mean heterozygosity within the same level, e.g. difference between ‘Non-genic’ and ‘Genic’ classes on the first level or difference between ‘Non-synonymous’ and ‘Synonymous’ classes on the third level.

### Patterns of genetic diversity in different genes

We have investigated the patterns of genetic diversity in those 6 303 chicken genes, for which at least 10 SNPs were mapped to the gene, in comparison to the overall genetic diversity pattern. In particular, we wanted to find out if the decrease in genetic diversity is faster or slower in certain genes. The results for all the 6 303 genes are presented in S2 Table including the R^2^ and slope values. Reliabilities (R^2^) of the linear regression of the genetic distance from the wild ancestor on heterozygosity for the genes ranged from 0.036 to 0.701 with a mean R^2^ of 0.450 and the slopes ranged from −0.110 to −1.099. However, the R^2^ values were correlated to the number of SNPs within the genes with r = 0.562. The slopes were independent of the SNP numbers within genes with r = 0.026. The correlation between the slopes and R^2^ values was −0.556. We evaluated the regression coefficients (slopes) of the relationship between the heterozygosity and genetic distance for the genes in the top and lowest 5% ranges, which were in total 32 genes at each end. Based on these slope classifications, functional annotations of the genes were done for the combination of molecular function, biological and immune system processes as well as KEGG pathways using the ClueGo package. Based on the ClueGo results, none of the genes in the top 5% range formed any functional clusters while 4 of the genes (namely: EGFR, PAFAH1B1, PTPRS and RTN4) in the lowest 5% were associated with brain development.

Genes in the lowest 5% had slopes ranging from −0.110 to −0.319 while the top 5% ranged from −0.960 to −1.099 (S3 Table). The genes in the top 5% indicate rapid changes in genetic diversity due to the genetic distance of the chicken breeds to *G. gallus* while those in the lowest 5% indicate genetic diversity changes at a very slow rate in relation to the genetic distance. We obtained the individual gene functions for these genes in the lowest and top ranges from DAVID annotation platform (S3 Table). The figures showing the relationship between genetic diversity and genetic distance in these genes are shown in S1 File and S2 File for the top and lowest 5% ranges, respectively.

The genes in the top 5% slope range were associated with transmembrane transport (SLC25A6, SLC22A15, SLC4A3), protein transport and protein metabolic processes (SLMO1, ERO1L, UCHL5, KCNB1, CSE1L), and lipid metabolic processes (PLCXD1, MIR33, HADHA) among other functions. The transmembrane transport refers to the transportation of solute/s across the protein embedded lipid bilayer. The lipid bilayer facilitates the distribution of molecules such as ions and proteins between different membrane compartments by allowing them to cross to different areas only when it is necessary [25]. Proteins are responsible to perform a wide range of important biochemical functions including those relating to adaptation, survival and performance. Proteins and lipids are also core biological molecules of living organisms and key molecules for energy generation. The energy and nutrient requirements differ for different types of breeds or strains and are as well influenced by other factors such as breeding goals and management systems [26, 27]. Hence the high flexibility of these genes to change may also be associated with such factors in addition to the change in genetic diversity which was initially due to the populations’ physical expansion from the *G. gallus*. In general, these genes are flexible to change without necessarily causing harm to the individuals but probably to complement the evolution of the populations. The genes in this range had R^2^ ranging from 0.419 to 0.628 indicating the good association of the genetic diversity and the genetic distance to the wild populations.

Most of the genes in the lowest 5% slope range have consistently lower genetic diversity across the breeds despite the genetic distance to the *Gallus gallus* (see S2 File) and they are mainly related to critical functions which may be absolutely necessary for normal functioning of the individuals. Among all the genes, the slopes were the lowest and much closer to zero for the DPYSL2 (−0.112) and GRB2 (−0.110) genes which also had the lowest R^2^ values of 0.036 and 0.038, respectively among all the genes. The GRB2 gene, which is involved in many pathways and functional processes, is assumed to be highly conserved in chicken as well as in humans and was reported to be under very strong evolutionary constraint [28]. Other than some of the genes, which are mentioned above for being related to the development of the brain, genes in the lowest 5% range were also found to be associated with other important developmental processes, functions and pathways. Such include positive regulation of cell proliferation (NTF3, ESRP2, EGFR, FGFR1), positive regulation of reactive oxygen species metabolic process (GRB2, STK17A), regulation of cell death, cell and structure morphogenesis (GRB2, NTF3, DOCK5, EGFR, STK17A), positive regulation of reproduction (GNRH1), development of spinal cord (PTPRS), salivary gland morphogenesis (FGFR1, ESRP2, EGFR), lung morphogenesis (FGFR1, ESRP2), brain morphogenesis and development (FGFR1, PAFAH1B1, DPYSL2), axon development (NEFM, RAB8A, RTN4, DPYSL2) among others functions. ADAM28 belongs to the family of ADAMs genes, being a family of transmembrane proteins involved in several processes including embryonic morphogenesis and tissue development, neurogenesis, cell adhesion, cell migration, axon outgrowth and guidance, cell proliferation and cell differentiation during development [29]. In humans, the ADAMs are said to be involved in the regulation of growth factor activities, promoting cell growth and invasion. They may alter cell communication or signaling in cancer cells causing an increase in cancer cell proliferation and progression [30]. The allele frequency in our study showed a very rapid fixation of the alternative allele in the ADAM28 in all breed categories supporting the assumption that the mutations might be of importance. In general, the consistent lower genetic diversity in the lowest 5% slope range and limited/lack of response to the changes in genetic distance to *G. gallus* can be due to several reasons such as i) some genes may be under evolutionary constraints such that changes of the genes may be generally critical for normal development or functioning of the animal and changes in the genes may have detrimental effects. ii) Purifying selection may be acting to remove the non-favorable alleles and is, therefore, leading to rapid fixation of the other allele. iii) On the other hand, genetic diversity might have been already reduced from the founders i.e. selection and fixation of the preferred variants took place prior to domestication; hence no or less feasibility for further reduction in genetic diversity is being possible. In this line, we investigated the genes which have the lowest estimated heterozygosity within the *Gallus gallus* populations. We found out that 27 of the 32 genes in the lowest 5% slope range were among genes with the lowest 1% of estimated heterozygosity within *Gallus gallus*. Furthermore, seven of those 27 genes also were among the genes with the lowest 5% of estimated heterozygosity within all breed categories.

We have analyzed the patterns of genetic diversity within a wide range of chicken breeds as a function of genetic distances from the chicken wild types. Given all forces taking place in the genome, we can conclude that the overall genetic diversity in the chicken can be well explained by the genetic distance to the wild populations. However, different functional genomic regions, genes and pathways have shown different evolutionary dynamics across the breeds resulting in different patterns of the genetic diversity compared to the overall genome and the neutral loci. The non-synonymous sites in particular have shown to be the most deviating from the overall pattern of genetic diversity compared to other genomic sites. Furthermore, we have found that genetic diversity changed at a faster rate in genes which are flexible to be manipulated according to the population needs e.g. genes involved in energy metabolism. On the other hand, genes which show resistance to change are associated with critical vital functions e.g. brain development, crucial for normal functioning of the individuals. Such genes presumably have maintained similar low levels of genetic diversity across all populations by selection or by evolutionary constraints, and the variations or the lack thereof in the genomic diversity between the breeds (within these genes) does not reflect the genetic distances to the wild type populations. This study presents insights and contributes to the knowledge of evolutionary dynamics of different functional genomic regions in the chicken.

## Materials and Methods

### Ethics statement

The data used was derived from a previous study [7], sourced from the SYNBREED (http://www.synbreed.tum.de/) project which was funded by the German Federal Ministry of Education and Research (FKZ 0315528E). Sampling of chickens followed the German Animal Welfare regulations, the authorities of Lower Saxony were notified according to §8 of the German Animal Welfare Act (33.9-42502-05-10A064) and with the written consent of the animal owners.

### Data description and quality control

Data consisted of 3 235 chicken individuals from 174 chicken populations collected in Asia, Africa, South America and Europe. The populations were classified into twelve breed categories which were based on their continent of origin and/or type as described in S1 Table. The chickens were genotyped with the 600K Affymetrix® Axiom™ Genome-Wide Chicken Genotyping Array [31]. We used only the SNPs from the 28 autosomal chromosomes and removed 499 SNPs with ambiguous chromosome annotation. The data was filtered for an animal call rate of ≥95% and SNP call rate of ≥99% using the SNP & Variation Suite (SVS) version 8.1 [32]. We performed LD based pruning to account for ascertainment bias [33] using the PLINK software v1.9 [34, 35] with the parameters *indep 50 5 2*. After the filtering steps 156 753 SNPs were left for further analysis and imputation was performed to recover missing genotypes using Beagle 3.3 [36]. A further description of the data can be found in Malomane et al. [7].

### Classification of the SNPs

We classified SNPs according to their functional consequences and assigned them to their associated genes using the Affymetrix Galgal5 annotation map [37]. SNPs were classified into the following categories: non-synonymous which is made of the missense and nonsense (only eight in total) variants, synonymous, exonic (a combination of the non-synonymous and synonymous SNPs as well as other coding and non-coding exonic SNPs which were not assigned as non-synonymous or synonymous), intronic, 5’ untranslated region (5’ UTR), 3’ untranslated region (3’ UTR), upstream, downstream and intergenic classes. SNPs assignments were prioritized in the order as they appear on Table 1. For example, if one SNP is associated with two genes but has different functional consequences for the two genes (e.g. non-synonymous for one gene and synonymous for the other gene) then a non-synonymous functional consequence was considered first instead of the other consequences, followed by synonymous and so forth. As for the up- and downstream variants, a SNP was assigned to the upstream class if it was located within 5 kb upstream of the gene and in analogy for the downstream SNPs. The distribution of SNPs into their functional classes is shown in column 1 of Table 1.

For assigning SNPs to indiviadual genes, the 156K SNPs were mapped to a total of 10 456 associated genes [37].

### Estimation of genetic diversity outward from wild populations

Two subspecies of the wild populations (*Gallus gallus*), the *G. gallus spadiceus* and *G. gallus gallus*, sampled about 20 years ago were used as reference for original founders, and reflect genetic diversity in centers of domestication.

We estimated the pairwise Reynolds’ genetic distances [19] between the two wild type populations (*G. gallus* ssp.) and the domesticated populations, and then calculated the mean genetic distance of each domesticated population to the two wild populations. Furthermore, observed heterozygosity was estimated within each population. Then, we estimated the linear relationship between the overall genetic diversity within the domesticated populations and their mean genetic distances to the two wild type populations. The amount of variation in genetic diversity within the populations which can be explained by the genetic distance was measured by the R^2^ value. To investigate if different SNP classes and genes show similar patterns as the overall genome pattern (when using all SNPs), we also estimated the genetic diversity in the different SNP classes and in genes and subsequently estimated the linear relationship with the genetic distances to the wild populations. We used the likelihood ratio test implemented in the R lmtest package (v0.9-36) [38] which uses the the *χ*^2^ test to compare the linear regression coefficients of the overall pattern to the patterns of the different SNP classes.

For the individual genes, because some of the genes were annotated with only one or very few associated SNPs while others were annotated with more, we only considered genes with at least ten associated SNPs (resulting in 6 303 in total) for making comparisons with the overall pattern. We evaluated the rate of change in the genetic diversity within the genes due to the change in genetic distances of populations to the wild populations using the regression coefficients of the linear relationship between the two parameters.

### Functional annotation of genes

Genes within the lowest and highest 5% ranges of regression coefficients in the relationship between genetic diversity within populations and genetic distances to the wild populations were grouped into functional terms using the ClueGO (v2.5.1) [39] ontology enrichment package in Cytoscape (v3.6.1) [40]. Additionally, individual gene functions were annotated using the DAVID functional annotation tool (v6.8) [41].

## Acknowledgements

We acknowledge the members of the SCDP consortium as well as the breeders of the “Bund Deutscher Rassegeflügelzüchter e.V.” and the “Gesellschaft zur Erhaltung alter und gefährdeter Haustierrassen e.V” in Germany for providing samples or SNP data, or gave access to their animals for sampling.

## Supporting information

**S1 Table. Categories of the chicken breeds.** (DOCX)

**S2 Table. The R**^**2**^ **and slope values of the relationship between genetic diversity and genetic distances of populations to *Gallus gallus* ssp. estimated from the 6 303 genes.** (XLSX)

**S3 Table. List and functions of the genes in the top and lowest 5% slope ranges.** (DOCX)

**S1 Fig. Genetic diversity vs. Reynolds’ genetic distance to the *Gallus gallus* estimated from 1000 SNP samples in 100 replicates.** The dashed lines represent the 100 sample sets and the gray area shows a 95% confidence interval. (TIF)

**S2 Fig. The relationship between the observed heterozygosity and genetic differentiation (***F*_*ST*_**) from *G. gallus* (left), and the relationship between** *F*_*ST*_ **and Reynolds’ genetic distances to *G. gallus* (right).** The regression lines of the relationships are drawn in red. The R^2^ of the left figure is 0.862 and 0.990 for the right figure. Different breed categories are denoted in different colors and/or shapes. (TIF)

**S3 Fig. Comparison of the relationship between the genetic distances to *G. g*allus and the genetic diversity estimated from the non-synonymous class vs. 100 random samples of the same number of SNPs as the non-synonymous class from the overall SNPs**. The black dotted lines represent estimations with the overall SNPs, the red solid line represents the non-synonymous SNPs. The shaded areas represent the 95% confidence intervals of the regression lines. The mean R^2^ of the 100 samples is 0.869 and the mean slope is −0.684. (TIF)

**S4 Fig. Comparison of the relationship between the genetic distances to *G. gallus* and the observed heterozygosity estimated from intronic SNPs vs. the overall set.** The black dashed lines represent estimations with the 100 replicates from randomly sampling 1000 SNPs from the intronic SNPs and the red solid line represents overall SNPs. The 95% confidence intervals are shaded in gray. The mean R^2^ and slope of the 100 samples are 0.869 and −0.686, respectively. (TIF)

**S5 Fig. Comparison of the relationship between the genetic distances to *G. gallus* and the observed heterozygosity estimated from intergenic SNPs vs. the overall set.** The black dashed lines represent estimations with the 100 replicates from randomly sampling 1000 SNPs from the intergenic SNPs and the red solid line represents overall SNPs. The 95% confidence intervals are shaded in gray. The mean R^2^ and slope of the 100 samples are 0.865 and −0.678, respectively. (TIF)

**S1 File.** Figures showing the relationship between genetic diversity and genetic distance to *G. gallus* for genes in the top 5% slope range. (ZIP file containing TIF figures)

**S2 File.** Figures showing the relationship between genetic diversity and genetic distance to *G. gallus* for genes in the lowest 5% slope range. (ZIP file containing TIF figures)

